# Temporal and Age-Dependent Regulation of Phagocytosis-Related Signatures After Ischemic Stroke: *Cross-Species Transcriptomic Evidence*

**DOI:** 10.64898/2026.08.07.743522

**Authors:** Rami A. Shahror, Carol A. Morris, Mohamed A. Sadek, Esraa Shosha, Abdelrahman Y. Fouda

**Affiliations:** Department of Pharmacology & Toxicology, College of Medicine, University of Arkansas for Medical Sciences, Little Rock, AR, USA; Departments of Clinical Pharmacy and Pharmacology & Toxicology, Faculty of Pharmacy, Cairo University, Cairo, Egypt

**Keywords:** Ischemic stroke, efferocytosis, macrophages, aging, neuroinflammation, transcriptomics

## Abstract

**Background:** Efferocytosis, the phagocytic clearance of apoptotic and damaged cells, promotes inflammation resolution and tissue repair following ischemic stroke. This study investigated temporal changes in efferocytosis and phagocytosis-related transcriptional programs during acute experimental stroke, examined the effects of aging on these responses, and assessed whether similar immune signatures are present in human ischemic stroke.

**Methods:** Publicly available transcriptomic datasets from murine transient middle cerebral artery occlusion (tMCAO; GSE104036 and GSE112348), permanent middle cerebral artery occlusion (pMCAO; GSE137482), and human peripheral blood after ischemic stroke (GSE16561) were analyzed using OmicSoft/Ingenuity-style pathway analysis. Functional validation included in vivo assessment of efferocytosis after tMCAO and in vitro phagocytosis assays using bone marrow-derived macrophages from young and aged mice.

**Results:** Both acute tMCAO models exhibited robust inflammatory activation together with sustained activation of phagocyte-related pathways during the first 24 hours after stroke. Human peripheral blood demonstrated similar inflammatory and phagocytic signatures, supporting translational relevance. Increased efferocytosis at 24 hours after tMCAO was associated with neuroprotection. Although both young and aged mice activated phagocytosis-related pathways after pMCAO, aged mice showed reduced phagosome formation. Consistent with these findings, macrophages from aged mice exhibited enhanced inflammatory responses and impaired uptake of apoptotic cells.

**Conclusions:** A conserved post-stroke immune response characterized by inflammatory activation and phagocyte-mediated clearance was identified across murine and human datasets. Efficient efferocytosis was associated with neuroprotection, whereas aging impaired apoptotic cell clearance and promoted a pro-inflammatory macrophage phenotype, highlighting efferocytosis as a potential therapeutic target for ischemic stroke.

## Introduction

Ischemic stroke causes a complex cascade of neuronal injury, vascular stress, innate immune activation and cellular debris accumulation. The acute phase of stroke is critical because the inflammatory and clearance mechanisms activated during this window influence lesion expansion, and tissue remodeling.**^1^** Efferocytosis is the clearance of apoptotic cells and is emerging as an important mechanism that can shape the inflammatory environment after central nervous system injury.^2^ Recent studies have suggested that aging can reduce efferocytc efficiency and may be a therapeutic prospect for treating age-related diseases.^3, 4^

The premise of this study is that efferocytosis/phagocytosis is a protective process after stroke and that its activation may vary over time, with stroke severity, and with biological aging as recently reviewed by us.^2^ Post-stroke dynamic immune response evolves from an inflammatory activation early on that later shifts to a reparative clearance process.^5, 6^ A deeper understanding of when and how efferocytosis-related pathways are activated may help reveal time windows in which clearance is most efficient and beneficial for healing.^2^

Aging is the single strongest non-modifiable risk factor for ischemic stroke and age largely influences stroke outcomes. The risk of stroke incidence doubles every 10 years after age 55, and nearly three-quarters of all strokes occur in adults age 65 and older.^7, 8^ Furthermore, these populations often experience poorer stroke outcomes, higher mortality rates, long-term disability, and longer hospital stays compared to younger ischemic stroke patients.^9^

The National Institutes of Health (NIH) recent initiatives focus on data sharing and reuse, and in silico analysis in an effort to reduce animal testing while improving reproducibility and translational relevance.^10^ Therefore, recent studies have emphasized the need to reanalyze and utilize publicly available data to uncover new discoveries.^11-13^ Here, we integrated publicly available datasets spanning from acute tMCAO in young mice to age-dependent pMCAO in young and aged mice. Furthermore, we analyzed peripheral blood data from human ischemic stroke patients. This cross-dataset design enabled comparison of efferocytosis/phagocytosis-related pathway signatures across time after stroke, ischemic injury paradigms, biological aging, and species. Because transcriptomic pathway outputs cannot directly distinguish beneficial efferocytosis from broader phagocytic activity or phagoptosis, we interpret these signatures as evidence of phagocyte-associated clearance programs and place them in the context of complementary efferocytosis assays conducted in our laboratory.

## Methods

### Study Design and Data Sources

This study used transcriptomic outputs generated from publicly available mouse and human stroke datasets analyzed through QIAGEN Ingenuity Pathway Analysis (IPA), OmicSoft, and Interpret bioinformatics platforms:

https://omicsoft-explorer.ingenuity.com/

https://analysis.ingenuity.com/ipw/

### OmicSoft/IPA visualization and pathway analysis

For temporal analysis after transient focal ischemia, two wild-type (WT) mouse temporary middle cerebral artery occlusion (tMCAO) datasets were examined: GSE104036, corresponding to a 35-minute occlusion model at 6, 12, and 24 hours after tMCAO; and GSE112348, corresponding to a 60-minute occlusion model at the same acute time points. For age-related analysis, GSE137482 was used to compare young (3-month-old) and aged (18-month-old) WT mice at day 3 after permanent MCAO (pMCAO). For translational comparison with human stroke, GSE16561 was used to analyze human peripheral blood data at day 1 after ischemic stroke. Table 1 provides an overview of the datasets, including the figure numbers, sample sizes, and other details.

**Table 1:**
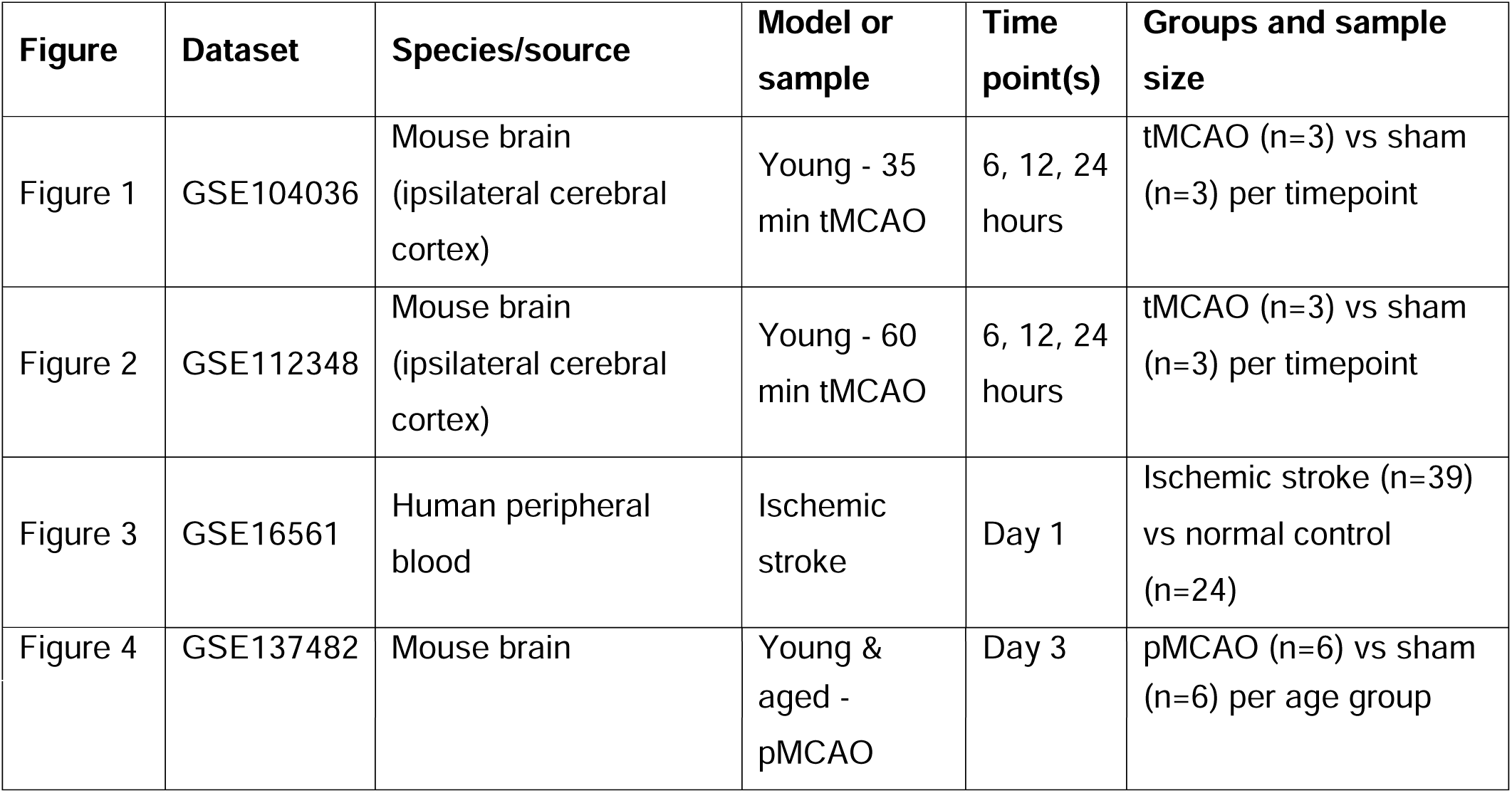
Dataset Overview.

### Transcriptomic and Pathway Analyses

For each dataset, we present volcano plots, graphical summaries, upstream regulator analyses, and outputs from canonical pathway/function analyses. The principal biological focus was on efferocytosis- and phagocytosis-related outputs. The canonical pathway used to represent phagocytosis was phagosome formation, whereas functional analyses included recruitment of phagocytes, activation of phagocytes, and phagocytosis of cells. A list of differentially expressed genes for each comparison is included in the supplementary Excel file 1.

### Middle cerebral artery occlusion model

Male C57BL/6J WT mice (12–16 weeks old) were subjected to right tMCAO for 1 hour using the intraluminal filament technique (Doccol 602145), followed by reperfusion as described by us.^14^ Body temperature was maintained at 37°C with a heating pad. Mice were sacrificed at 24 hours via trans-cardiac perfusion with phosphate-buffered saline (PBS); brains were collected and fixed in 4% paraformaldehyde (PFA) overnight at 4°C. Brains were then cryopreserved in sucrose solutions and prepared for cryosectioning by embedding in Optimal Cutting Temperature (O.C.T).

### *In vivo* efferocytosis assay

To investigate efferocytosis in the infarct region of the mouse brain, five 10-μm coronal sections were obtained from fixed brain tissue embedded in O.C.T. via cryosectioning. These sections underwent immunofluorescent labeling as previously described.^15^ Efferocytosis was assessed by co-localizing Iba-1 positive microglia/macrophages (MΦ) with apoptotic cells identified by TUNEL (Terminal deoxynucleotidyl transferase dUTP nick end labeling) in the peri-infarct area. The analysis was conducted across five equally spaced brain coronal sections that encompassed the middle cerebral artery (MCA) territory. For each section, five high-power z-stack images were captured from the peri-infarct area. The efferocytosis index, defined as the percentage of dead cells engulfed by microglia/MΦ, was calculated using the following equation: [number of Iba1^+^TUNEL^+^ cells / number of TUNEL^+^ cells] × 100%.

### *In vitro* efferocytosis assay

The *in vitro* efferocytosis assay was conducted as we previously described.^16^ Non-apoptotic (NACs) or apoptotic (ACs) K562 suspension culture cells, labeled with Dil red-tracker, were added to MΦ labeled with CFDA-green tracker, in a 5:1 ratio and incubated for 45 minutes to allow for the efferocytosis of ACs by MΦ. After incubation, the residual unengulfed suspension K562 cells were removed by washing the culture three times with PBS. The MΦ culture with internalized ACs was then fixed and processed for confocal imaging. Efferocytosis was determined as the percentage of MΦ engaged in efferocytosis using the following formula: in vitro efferocytosis (%) = [number of MΦ internalized ACs ÷ total number of MΦ] × 100.

#### Macrophage isolation and LPS stimulation

Bone marrow-derived MΦ were isolated from 2 and 15-month-old WT mice and stimulated with lipopolysaccharide (LPS) at a concentration of 100 ng/mL, or treated with vehicle, for 18 hours as described previously to elicit an inflammatory response.^17^ Western blotting on cell lysate using anti-TNFα (Abcam, Cat. # ab1793) was conducted as previously described.^17^

#### Data Presentations and Statistical Analysis

For the analyzed datasets, volcano plots, differentially expressed genes (DEGs), P-values, and activation z-scores were generated using Qiagen OmicSoft Explorer and IPA Interpret bioinformatics software. For experiments conducted in our lab, the significance of differences between 2 groups was determined by Student’s t-test, with P<0.05 considered statistically significant. Graphs and statistics were conducted using GraphPad Prism 11.

## Results

### Acute 35-Minute tMCAO Shows Early Activation of Inflammatory and Phagocytic/Efferocytosis-Related Programs

The 35-minute tMCAO dataset, GSE104036, was analyzed at 6, 12, and 24 hours after stroke. Figure 1 presents volcano plots for each time point, graphical summaries, upstream regulators, and canonical pathway/function outputs. Across the acute time window, volcano plots show robust upregulation of several genes with a lesser number of genes being downregulated (Figure 1A-C). Graphical summaries of the differentially expressed genes (DEGs) show a robust inflammatory response across the examined timepoints (Figure 1D-F). This was confirmed by the upstream regulators’ analyses which showed robust activation of inflammatory pathways, including Tumor Necrosis Factor-alpha (TNF-α), Toll-like Receptor 4 (TLR4) activation (represented by lipopolysaccharide - LPS), Interleukin-1 beta (IL-1β), and Interferon gamma (IFNγ) (Figure 1G-I).

**Figure 1.**
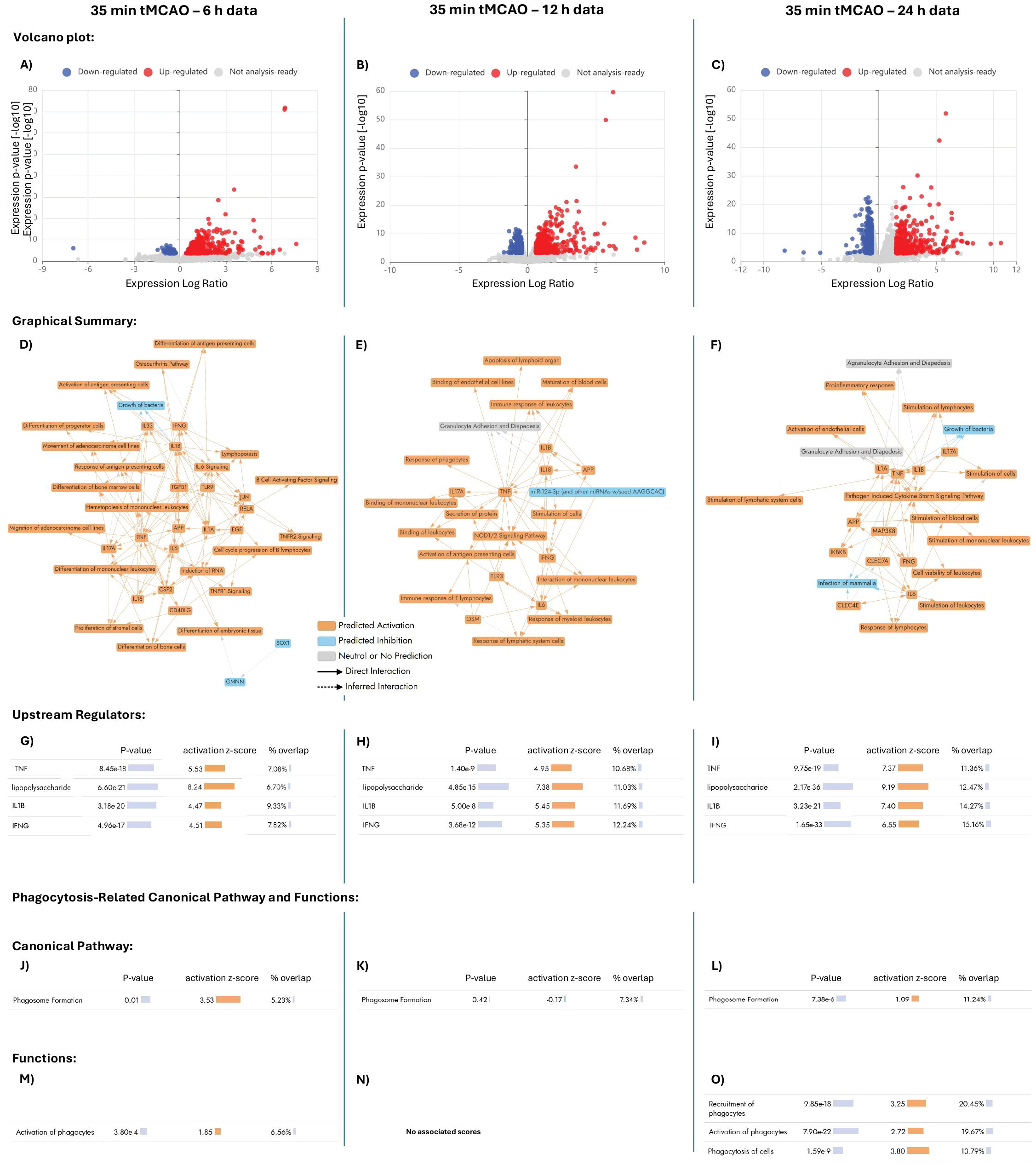
Acute transcriptional and pathway responses after 35-minute tMCAO in mice. **A-O)** Data from GSE104036 were analyzed at 6, 12, and 24 hours after 35-minute transient middle cerebral artery occlusion. Panels include volcano plots, graphical summaries, upstream regulator analyses, and canonical pathway/function outputs. Across acute time points, pathway/function outputs include phagosome formation, activation of phagocytes, recruitment of phagocytes, and phagocytosis of cells.

Canonical pathway/function outputs support an early engagement of phagocyte-associated clearance programs within 6 h of injury with phagosome formation and activation of phagocytes showing positive scores (Figure 1J, M). The 12-hour time point, however, does not show positive phagocytic activation scores (Figure 1K, N). Yet, the 24-hour timepoint shows robust phagocytic activation, as evidenced by positive z-scores for all phagocytic parameters: phagosome formation, recruitment of phagocytes, activation of phagocytes, and phagocytosis of cells (Figure 1L, O). Collectively, these findings suggest that a short transient occlusion model engages early immune and clearance-related transcriptional responses within hours of reperfusion with a possible biphasic phagocytic response.

### A 60-Minute tMCAO Model Shows Acute Activation of Phagocyte-Related Pathways Across 6, 12, and 24 Hours

The 60-minute tMCAO dataset, GSE112348, was analyzed at 6, 12, and 24 hours after stroke. Figure 2 includes matched volcano plots, graphical summaries, upstream regulator analyses, and canonical pathway/function outputs for the three acute time points. As in the 35-minute model, the 60-minute tMCAO dataset shows strong upregulation of DEGs transcripts with a strong inflammatory signature that included TNF-α, IL-1β, and IFNγ (Figure 2A-I). Similar to the 35-minute model, the 60-minute tMCAO dataset shows recurring phagocyte activation and clearance-related pathway categories, including phagosome formation, recruitment of phagocytes, activation of phagocytes, and phagocytosis of cells. More specifically, the canonical pathway, phagosome formation, showed positive z-scores for 6 and 12 hours time points but not at 24 hours (Figure 2J-L). All phagocytic functional pathways showed positive z-scores at all time points (Figure 2M-O).

**Figure 2.**
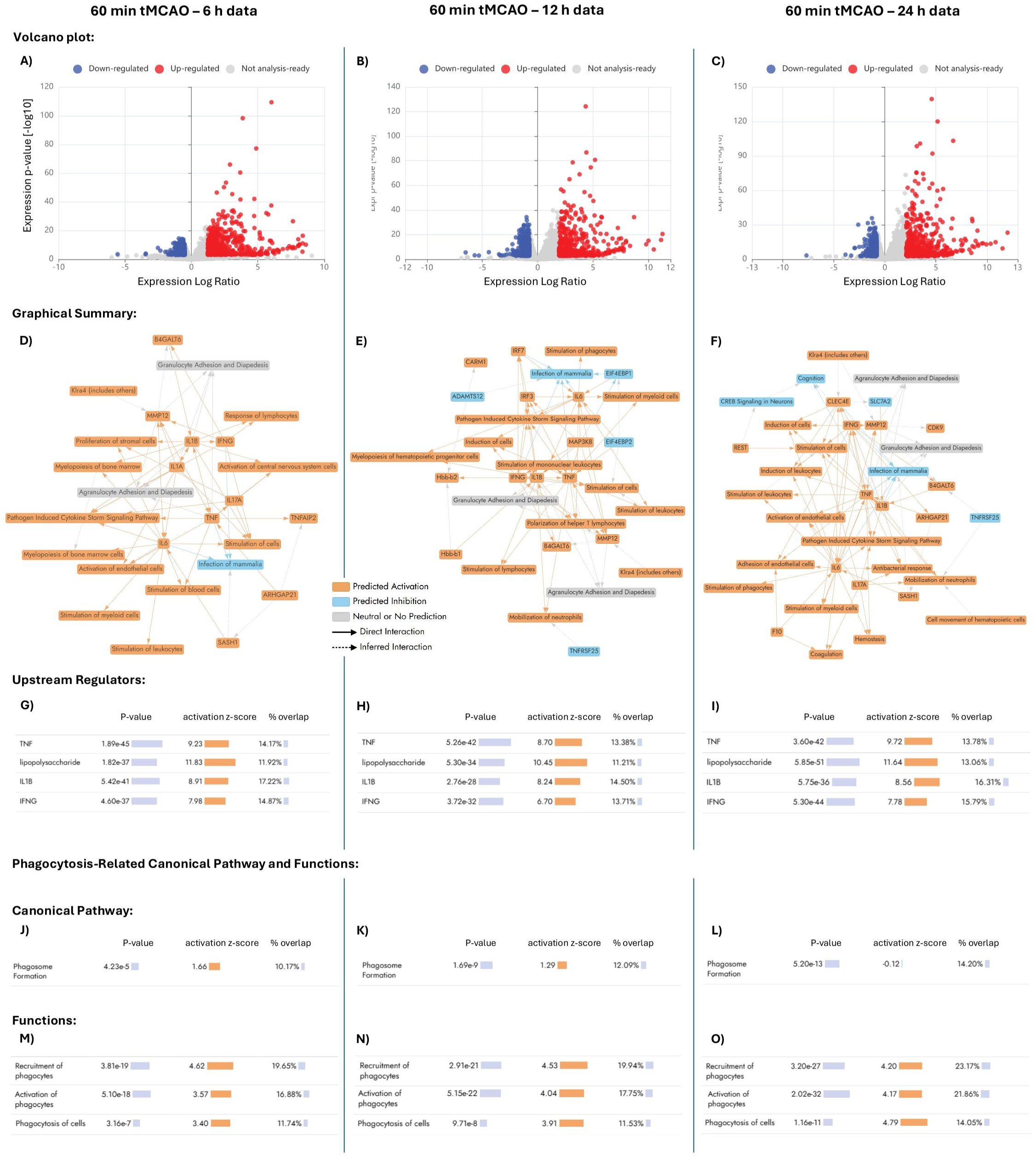
Acute transcriptional and pathway responses after 60-minute tMCAO in mice. **A-O)** Data from GSE112348 were analyzed at 6, 12, and 24 hours after 60-minute transient middle cerebral artery occlusion. Volcano plots, graphical summaries, upstream regulators, and canonical pathway/function outputs demonstrate acute inflammatory and phagocyte-associated pathway activation. Pathway/function outputs include phagosome formation, recruitment of phagocytes, activation of phagocytes, and phagocytosis of cells.

Collectively, these data show that both transient ischemia models exhibit convergent activation of upstream inflammatory regulators and phagocyte- and efferocytosis-related pathways during the first 24 hours after stroke.

### Human Peripheral Blood at Day 1 After Ischemic Stroke Shows Convergent Inflammatory and Phagocyte-Related Signatures

To assess whether the inflammatory and phagocytic signature observed in the murine stroke model is also present in clinical cases, we analyzed human peripheral blood data from GSE16561 collected on day 1 after ischemic stroke. This analysis included a volcano plot, a graphical summary, upstream regulators, and outputs related to canonical pathways and functions, as illustrated in Figure 3.

**Figure 3.**
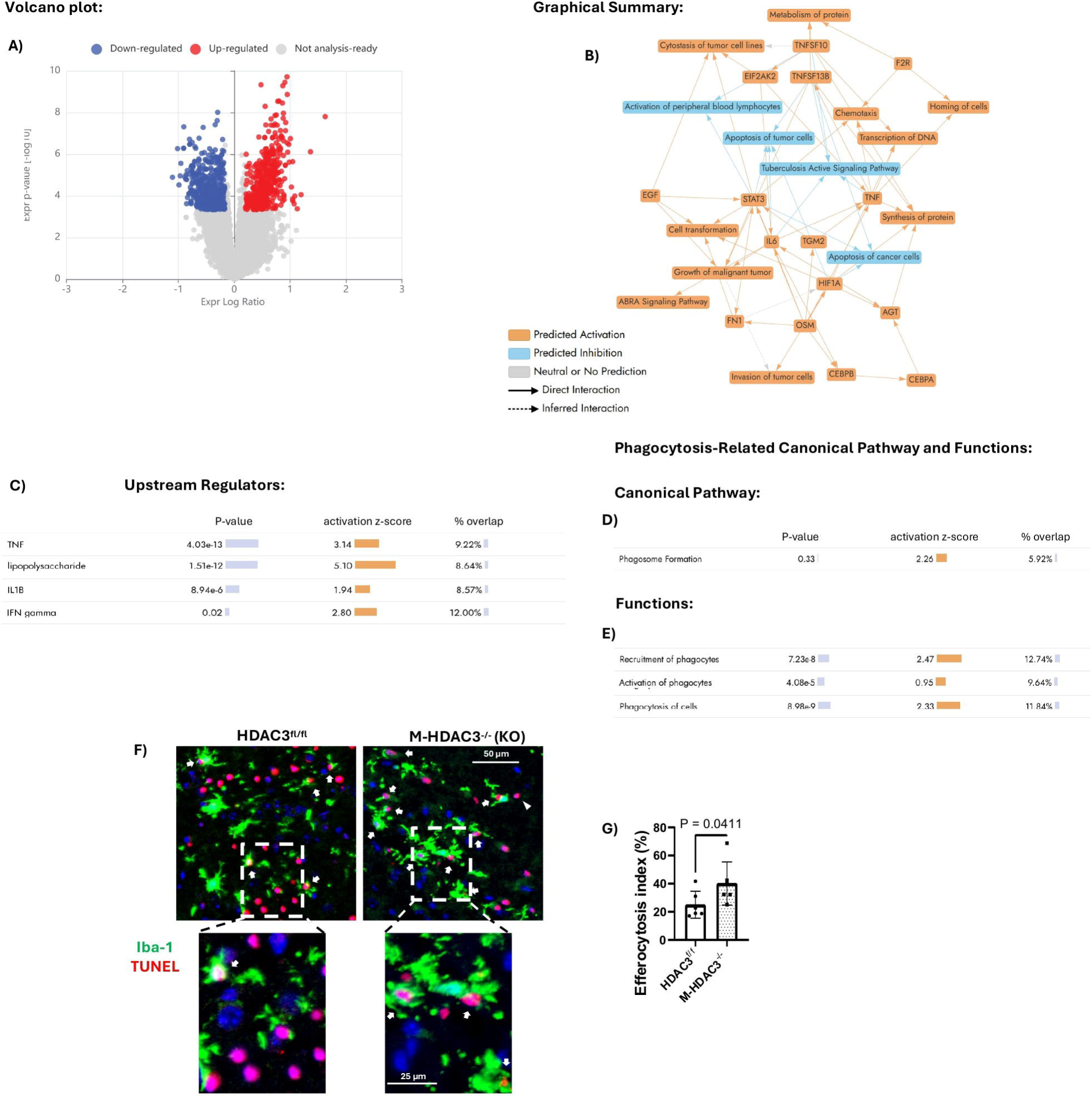
Human peripheral blood pathway signatures at day 1 after ischemic stroke and validation of brain efferocytosis at 24 hours after tMCAO. **A-E)** Human peripheral blood data from GSE16561 were analyzed at day 1 after ischemic stroke. Panels include volcano plot, graphical summary, upstream regulators, and canonical pathway/function outputs, including phagosome formation, recruitment of phagocytes, activation of phagocytes, and phagocytosis of cells. **F, G)** Representative images and quantification show Iba-1-positive microglia/MΦ colocalized with TUNEL-positive cellular material in floxed control and myeloid HDAC3 knockout mice at 24 hours after tMCAO, supporting enhanced efferocytic clearance in the myeloid HDAC3 knockout condition. Arrowheads represent free apoptotic cells, and arrows represent active efferocytosis.

The human dataset shows pathway/function categories aligned with the mouse datasets, including a strong inflammatory response, phagosome formation, recruitment of phagocytes, activation of phagocytes, and phagocytosis of cells, with all parameters showing positive z-scores (Figure 3A-E).

The detection of similar pathway themes in human blood supports the translational relevance of the mouse findings. However, because the human data derive from peripheral blood rather than brain tissue, the results should be interpreted as evidence of systemic post-stroke immune activation rather than direct proof of brain-resident efferocytosis.

To complement the pathway-based analysis with experimental evidence of efferocytosis, we assessed engulfment of TUNEL-positive ACs by Iba-1-positive microglia/MΦ in the mouse ischemic brain at 24 hours after tMCAO. This validation experiment compared myeloid histone deacetylase 3 (HDAC3) knockout mice with floxed controls, because recent reports support a protective role for myeloid HDAC3 deletion after ischemic stroke.^18, 19^ The representative images and quantification show increased colocalization of Iba-1-positive phagocytes with TUNEL-positive ACs in myeloid HDAC3 knockout mice, consistent with enhanced efferocytic clearance in a protected post-stroke context (Figure 3F, G). These data support the interpretation that the 24-hour phagocyte/efferocytosis-related transcriptomic signal corresponds, at least in part, to functional cellular clearance activity rather than pathway annotation alone.

### Young and Aged Mice both Show Phagocytosis-Related Pathway Activation at Day 3 after pMCAO

We next aimed to determine the effect of age on the post-stroke inflammatory and phagocytic responses. The age-comparison dataset, GSE137482, examined young 3-month-old and aged 18-month-old WT mice at day 3 after permanent MCAO (pMCAO). Figure 4 includes volcano plots for young and aged mice, corresponding graphical summaries, upstream regulator outputs, and canonical pathway/function outputs.

**Figure 4.**
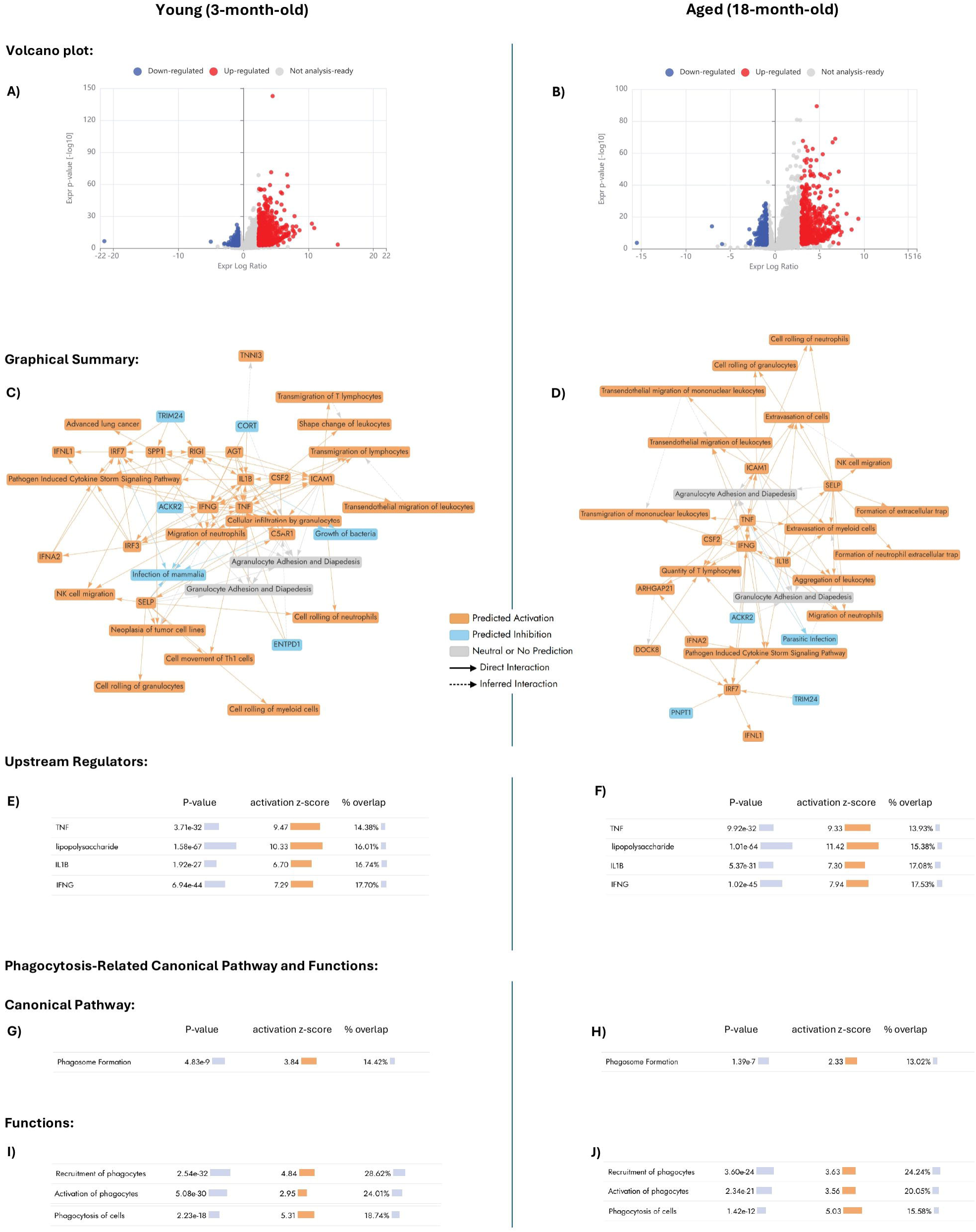
Age-associated pathway signatures at day 3 after pMCAO. **A-J)** Data from GSE137482 compare young 3-month-old and aged 18-month-old mice at day 3 after permanent MCAO. Volcano plots, graphical summaries, upstream regulator analyses, and canonical pathway/function outputs are shown for both age groups. Both young and aged mice show pathway/function outputs involving phagosome formation, recruitment of phagocytes, activation of phagocytes, and phagocytosis of cells; however, the aged group shows a relatively lower phagosome formation signal, suggesting possible age-associated impairment in clearance-related biology.

Both young and aged mice show strong inflammatory response (Figure 4A-F) and positive phagocytosis-related pathway/function outputs, including phagosome formation, recruitment of phagocytes, activation of phagocytes, and phagocytosis of cells (Figure 4G-J). Of note, the aged group showed a relatively lower phagosome formation z-score as compared to young mice, suggesting a possible phagocytic impairment with age (Figure 4G, H).

### Effect of age on macrophage inflammatory and phagocytic function

To further dissect the effect of aging on MΦ inflammatory and efferocytosis response. We isolated and compared bone marrow-derived MΦ from young and aged mice. In separate experiments, we stimulated MΦ with LPS to simulate the stroke inflammatory environment and also conducted an efferocytosis experiment as we previously described.^16, 17^ Aged MΦ displayed a heightened inflammatory response to LPS stimulation, as shown by greater TNF-α protein expression relative to MΦ from young mice (Figure 5A, B). In parallel, aged MΦ exhibited reduced engulfment of apoptotic cells in the *in vitro* efferocytosis assay (Figure 5C, D). Together, these findings suggest that aging shifts MΦ toward a more pro-inflammatory state while reducing their capacity to clear apoptotic targets, a combination that could impair inflammatory resolution after stroke despite the presence of phagocytosis-related pathway activation in aged post-stroke tissue.

**Figure 5.**
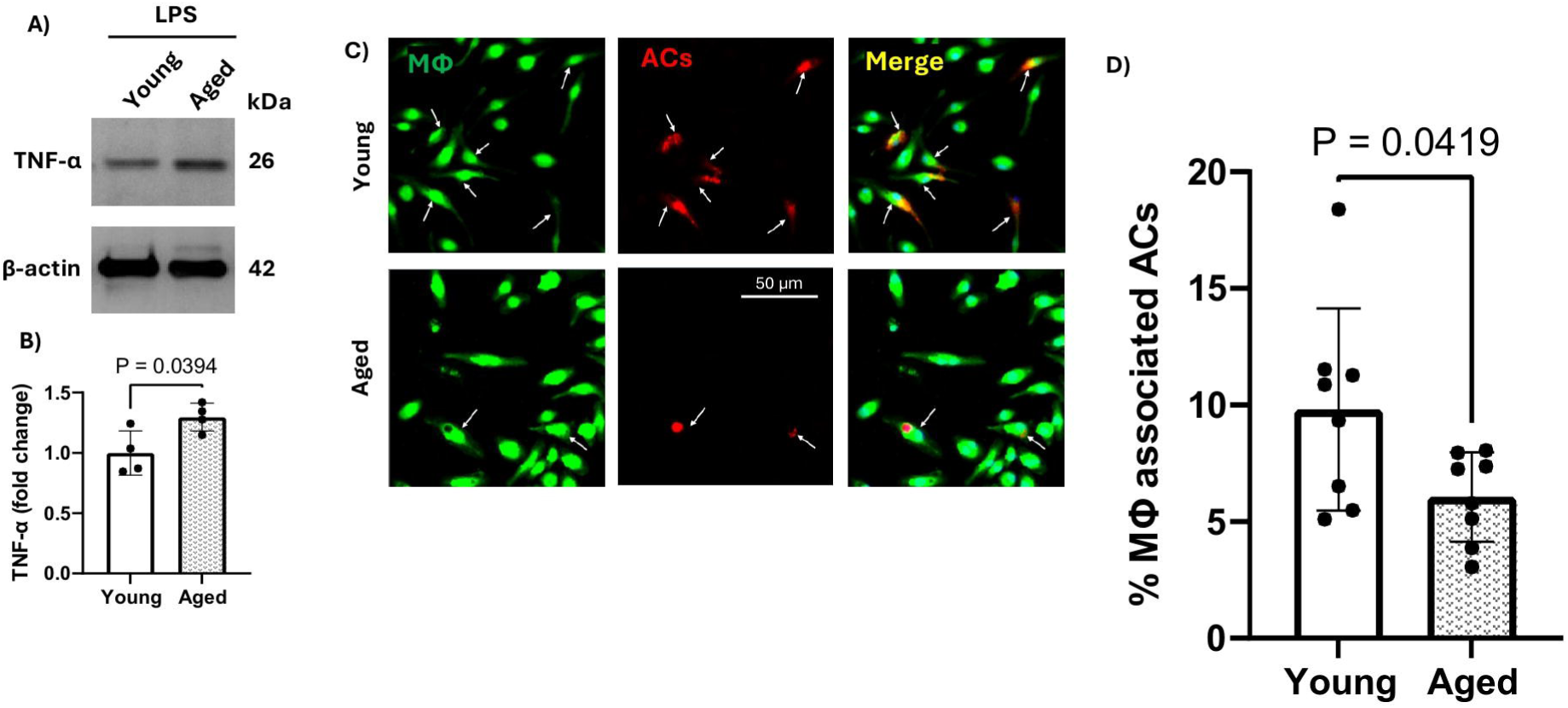
MΦ from aged mice show impaired efferocytosis *in vitro*. **A, B)** Bone marrow-derived macrophages (MΦ) from aged mice show increased TNF-α protein expression after LPS stimulation compared with MΦ from young mice, as determined by Western blotting and densitometric quantification. **C, D)** In vitro efferocytosis assay showing reduced uptake of apoptotic cells (ACs; CM-DiI red) by aged MΦ labeled with CFDA green tracker compared with young MΦ. Arrows indicate internalized ACs. Scale bar, 50 μm. n=8.

## Discussion

This study integrates transcriptomic pathway outputs from multiple mouse stroke datasets and a human ischemic stroke peripheral blood dataset to investigate the temporal, age-related, and translational features of efferocytosis/phagocytosis-associated responses after ischemic stroke. The major finding is that phagocyte-related pathways and functions are consistently activated across acute mouse tMCAO models, young and aged pMCAO mice, and patients with ischemic stroke, with the 24-hour timepoint showing a correlation between efferocytosis and positive stroke outcomes. We also found that aging can induce an amplified inflammatory response and reduce efferocytosis in macrophages. Collectively, these findings suggest that stroke rapidly induces immune programs associated with phagocyte activation, cellular debris clearance, and inflammatory regulation, and that aging may impair reparative efferocytosis.

The temporal analyses revealed that both the 35-minute and 60-minute tMCAO models exhibited robust activation of upstream inflammatory regulators, including TNF-α, IL-1β, IFNγ, and TLR-related signaling, as well as enrichment of phagosome formation, phagocyte recruitment, phagocyte activation, and phagocytosis-related functions. These observations support the concept that innate immune activation and clearance mechanisms are engaged early after ischemic injury and remain active throughout the acute post-stroke period. While excessive inflammation can contribute to secondary tissue injury, activation of phagocytes is also essential for the removal of apoptotic cells and cellular debris, suggesting a complex balance between detrimental and reparative immune responses.

The age-comparison dataset provides additional insight into how aging may influence post-stroke clearance mechanisms. Although both young and aged mice demonstrated activation of phagocytosis-related pathways, aged animals exhibited a relatively lower phagosome formation signal, suggesting a potential impairment in clearance efficiency. This interpretation is supported by our complementary validation studies showing that MΦ isolated from aged mice displayed enhanced TNF-α induction following LPS stimulation and reduced uptake of apoptotic cells during *in vitro* efferocytosis assays. Together, these findings suggest that aging does not abolish phagocyte activation after stroke; rather, it may shift myeloid cells toward a more pro-inflammatory phenotype while reducing their capacity to efficiently clear dying cells. Such an imbalance could contribute to prolonged inflammation, impaired tissue remodeling, and poorer functional recovery observed in older stroke patients.

The increased inflammatory response identified in our analysis underscores TNF-α’s crucial role across all examined datasets as both a central molecule and an upstream regulator. Previous studies have shown that TNF-α can induce apoptosis after a stroke, while other research has found that it can hinder the process of efferocytosis in various disease conditions.^20-23^ This dual mechanism suggests that TNF-α has a harmful role by both causing cell death and preventing the clearance of dead cells. However, despite this dual function, there is currently no TNF-based therapy available for stroke, likely because of its dual and time-dependent role after stroke.^22^ Therefore, there is a need to identify TNF-α modulators that promote post-stroke repair. The IPA analysis predicted LPS, which is derived from the outer membrane of gram-negative bacteria, as an upstream regulator in these stroke datasets representing sterile inflammation. This is likely due to robust TLR4 activation following a stroke. This evidence suggests that using LPS as a model for *in vitro* stroke research offers a valuable approach, despite its bacterial origin.

The human dataset further strengthens the translational relevance of these findings. Similar inflammatory and phagocyte-associated signatures were detected in patients with ischemic stroke, suggesting that immune-clearance programs are conserved across species and can be detected systemically following cerebral ischemia. However, it is important to note that the human data is derived from peripheral blood and therefore represents systemic activation rather than brain tissue activation which is a translational limitation when comparing these datasets. Importantly, the inclusion of tissue- and cell-based validation experiments provides additional evidence that transcriptomic signatures correspond to biologically relevant processes. Enhanced efferocytosis observed in myeloid HDAC3 knockout mice, a model previously associated with improved stroke outcomes, supports the concept that efficient apoptotic cell clearance may contribute to neuroprotection, with the 24-hour time point optimal for assessing efferocytosis. Conversely, impaired efferocytosis in aged MΦ highlights that the quality and efficiency of phagocyte responses may differ with age and inflammatory state. Together, these findings suggest that activation of phagocyte-related pathways alone may not be sufficient to predict beneficial outcomes; rather, the effectiveness of efferocytic clearance is likely a critical determinant of inflammatory resolution and tissue recovery after stroke.

## Limitations

Several limitations should be considered when interpreting these findings. First, the human dataset was derived from peripheral blood collected one day after ischemic stroke and therefore may not directly reflect cellular interactions occurring within the ischemic brain. Second, efferocytosis and phagocytosis are related but distinct biological processes, and transcriptomic pathway analyses cannot definitively distinguish between beneficial apoptotic-cell clearance and other forms of phagocytosis. Furthermore, the available pathway analysis platforms did not include a canonical efferocytosis-specific pathway. Consequently, phagosome formation and related functional categories—including recruitment of phagocytes, activation of phagocytes, and phagocytosis of cells—were used as surrogate indicators of efferocytosis-associated biology. While these pathways support activation of cellular clearance programs, they cannot distinguish reparative efferocytosis from potentially detrimental processes such as phagoptosis of stressed but viable cells.^24^ Future studies incorporating cell-specific analyses, ligand-receptor interactions, dedicated efferocytosis markers, and functional engulfment assays will be necessary to establish the precise role of efferocytosis in post-stroke recovery and determine whether these responses are ultimately beneficial or maladaptive.

## Conclusion

Taken together, the data support a model in which ischemic stroke triggers early inflammatory signaling that coincides with activation of phagocyte recruitment, phagosome formation, and cellular clearance pathways. In the injured brain, these responses likely involve both resident microglia and infiltrating myeloid cells, whereas in peripheral blood they reflect systemic immune activation following cerebral ischemia. Although these processes may represent an adaptive attempt to remove damaged cells and promote tissue repair, their biological impact likely depends on the timing, efficiency, and resolution of the clearance response. The accompanying validation studies further suggest that enhanced efferocytosis may be associated with protective outcomes, whereas aging may impair effective apoptotic-cell clearance despite persistent activation of phagocyte-related pathways. Overall, these findings identify efferocytosis-associated immune clearance programs as conserved components of the early post-stroke response and highlight their potential as therapeutic targets for promoting inflammatory resolution and recovery after ischemic stroke.

## Supporting information

Supplemental file 1 Datasets differentially expressed gene

## ACKNOWLEDGEMENTS

Funding: This work was supported by the National Institutes of Health R01 EY035658 to AYF. American Heart Association Postdoctoral Fellowship (25POST1372631) to RAS, National Center for Advancing Translational Sciences T32 to CAM (TR004918), American Heart Association Career Development Award (25CDA1446415) to ES.

## DECLARATION OF COMPETING INTERESTS

The authors declare that they have no competing financial interests or personal relationships that could have influenced the work reported in this paper.

### Declaration of generative AI and AI-assisted technologies in the manuscript preparation process

Qiagen IPA uses machine learning for analysis and generation of graphical summaries. Microsoft Copilot was used by the authors for literature search and language editing. The authors generated, interpreted, reviewed, edited, and approved the final version of the manuscript and take full responsibility for the content.

## References

1. Iadecola C, Buckwalter MS, Anrather J. Immune responses to stroke: mechanisms, modulation, and therapeutic potential. The Journal of clinical investigation 2020;130(6):2777–88.

2. Morris CA, Sadek MA, Modi P, Abdelnaem S, Rusch NJ, Shosha E et al. The role of efferocytosis in ischemic stroke and insights from retinopathy. Trends in neurosciences (Regular ed.) 2025;48(8):624–39.

3. Zhang M, Wei J, Sun Y, He C, Ma S, Pan X et al. The efferocytosis process in aging: Supporting evidence, mechanisms, and therapeutic prospects for age-related diseases. Journal of advanced research 2025;69:31–49.

4. Liu Z, Li Y, Ren Y, Chen J, Weng S, Zhou Z et al. Efferocytosis: The Janus-Faced Gatekeeper of Aging and Tumor Fate. Aging Cell 2025;24(2):e14467,n/a.

5. Zhang W, Zhao J, Wang R, Jiang M, Ye Q, Smith AD et al. Macrophages reprogram after ischemic stroke and promote efferocytosis and inflammation resolution in the mouse brain. CNS Neuroscience & Therapeutics 2019;25(12):1329–42.

6. Kim E, Cho S. Microglia and Monocyte-Derived Macrophages in Stroke. Neurotherapeutics 2016;13(4):702–18.

7. Howard G, Banach M, Kissela B, Cushman M, Muntner P, Judd SE et al. Age-Related Differences in the Role of Risk Factors for Ischemic Stroke. Neurology 2023;100(14):e1444–53.

8. Yousufuddin M, Young N. Aging and ischemic stroke. Aging (Albany, NY.) 2019;11(9):2542–4.

9. Lui SK, Nguyen MH. Elderly Stroke Rehabilitation: Overcoming the Complications and Its Associated Challenges. Current gerontology and geriatrics research 2018;2018(2018):1–9.

10. Anonymous. Strengthening Replication and Reproducibility of NIH-funded Research. National Institutes of Health 2026;

11. Ahmed M, Kim HJ, Kim DR. Maximizing the utility of public data. Frontiers in genetics 2023;14:1106631.

12. Sielemann K, Hafner A, Pucker B. The reuse of public datasets in the life sciences: potential risks and rewards. PeerJ 2020;8:e9954.

13. Pulley JM, Rhoads JP, Jerome RN, Challa AP, Erreger KB, Joly MM et al. Using What We Already Have: Uncovering New Drug Repurposing Strategies in Existing Omics Data. Annual Review of Pharmacology and Toxicology 2020;60(1):333–52.

14. Fouda AY, Eldahshan W, Xu Z, Lemtalsi T, Shosha E, Zaidi SA et al. Preclinical investigation of Pegylated arginase 1 as a treatment for retina and brain injury. Experimental neurology 2022;348:113923.

15. Fouda AY, Alhusban A, Ishrat T, Pillai B, Eldahshan W, Waller JL et al. Brain-Derived Neurotrophic Factor Knockdown Blocks the Angiogenic and Protective Effects of Angiotensin Modulation After Experimental Stroke. Mol Neurobiol 2017;54(1):661–70.

16. Shahror RA, Zaman B, Chuesiang P, Wild M, Shosha E, Leung Y et al. CD5L promotes efferocytosis and resolution of retinal ischemic injury. Cell Death Dis 2026;17(1):520.

17. Shosha E, Shahror RA, Morris CA, Xu Z, Lucas R, McGee-Lawrence ME et al. The arginase 1/ornithine decarboxylase pathway suppresses HDAC3 to ameliorate the myeloid cell inflammatory response: implications for retinal ischemic injury. Cell Death Dis 2023;14(9):621.

18. Zhang Y, Li J, Zhao Y, Huang Y, Shi Z, Wang H et al. Arresting the bad seed: HDAC3 regulates proliferation of different microglia after ischemic stroke. Science advances 2024;10(10):eade6900.

19. Li J, Wang C, Zhang Y, Huang Y, Shi Z, Zhang Y et al. Orchestrating the frontline: HDAC3-miKO recruits macrophage reinforcements for accelerated myelin debris clearance after stroke. Theranostics 2025;15(2):632–55.

20. McPhillips K, Janssen WJ, Ghosh M, Byrne A, Gardai S, Remigio L et al. TNF-α Inhibits Macrophage Clearance of Apoptotic Cells via Cytosolic Phospholipase A2 and Oxidant-Dependent Mechanisms. The Journal of immunology (1950) 2007;178(12):8117–26.

21. Borges VM, Vandivier RW, McPhillips KA, Kench JA, Morimoto K, Groshong SD et al. TNFα inhibits apoptotic cell clearance in the lung, exacerbating acute inflammation. American journal of physiology. Lung cellular and molecular physiology 2009;297(4):L586–95.

22. Kołodziejska R, Pawluk H, Tafelska-Kaczmarek A, Pawluk M, Koper K, Godlewski A et al. The Role of TNF-α in Ischemic Stroke. International journal of molecular sciences 2026;27(3):1424.

23. Kojima Y, Volkmer J, McKenna K, Civelek M, Lusis AJ, Miller CL et al. CD47-blocking antibodies restore phagocytosis and prevent atherosclerosis. Nature 2016;536(7614):86–90.

24. Brown GC. Neuronal Loss after Stroke Due to Microglial Phagocytosis of Stressed Neurons. International Journal of Molecular Sciences 2021;22(24):13442.

